# Elastin-like polypeptide delivery of anti-inflammatory peptides to the brain following ischemic stroke

**DOI:** 10.1101/2023.03.15.532834

**Authors:** John Aaron Howell, Nicholas Gaouette, Mariper Lopez, Stephen P. Burke, Eddie Perkins, Gene L. Bidwell

## Abstract

Inflammatory processes are activated following ischemic strokes and lead to increased tissue damage for weeks following the ischemic insult, but there are no approved therapies that target this inflammation-induced secondary injury. Here, we report that SynB1-ELP-p50i, a novel protein inhibitor of the nuclear factor kappa B (NF-κB) inflammatory cascade bound to drug carrier elastin-like polypeptide (ELP), is able to enter both neurons and microglia, cross the blood-brain barrier, localize exclusively in the ischemic core and penumbra in Wistar-Kyoto and spontaneously hypertensive rats (SHRs), and reduce infarct volume in male SHRs. Additionally, in male SHRs, SynB1-ELP-p50i treatment improves survival for 14 days following stroke with no effects of toxicity or peripheral organ dysfunction. These results show high potential for ELP-delivered biologics for therapy of ischemic stroke and other central nervous system disorders and further support targeting inflammation in ischemic stroke.

## Introduction

Stroke is the fifth leading cause of death in the United States ^1^. The majority (∼87 %) of strokes are ischemic and are caused by occlusion of a major cerebral artery ^1^. Ischemic strokes result in a lack of oxygen and energy reaching the tissue, the formation of reactive oxygen species, increases in glutamate release and intracellular calcium, and activation of inflammatory processes which can cause secondary tissue injury ^1–3^. Current treatments, tissue plasminogen activator (tPA) ^4^ or mechanical endovascular thrombectomy ^5^, can restore blood flow, but they do not treat the downstream molecular pathways that can worsen injury following stroke.

Inflammation is a major contributor to the tissue damage and neurological deficits seen following ischemic stroke ^3,6,7^. Beginning at stroke onset and continuing for days to weeks following the stroke, inflammatory signaling affects all cell types that comprise the neurovascular unit and can severely damage the remaining penumbral tissue surrounding the ischemic core ^3,8,9^. Following ischemic stroke, there is a marked increase in release of proinflammatory cytokines and reactive oxygen species that result in increased expression of adhesion molecules that allow for peripheral leukocytes to adhere and accumulate within the cerebrovasculature ^9,10^. These changes result in increases in activated microglia and astrocytes in the ischemic territory. When the blood-brain barrier is weakened by increased activity of matrix metalloproteinases, these peripheral leukocytes can migrate across the endothelium and cause further inflammation and tissue infarction ^3,10^.

Preclinical studies have shown that inhibiting inflammation both locally and systemically can reduce tissue damage and improve neurological outcome following ischemic stroke ^11–13^. However, these studies have not led to translation to the human population for a number of reasons, including lack of delivery of the therapeutic to the infarct and ischemic penumbra, the unknown and non-specific mechanism of action of many anti-inflammatory agents, and the unknown optimal treatment window following ischemia and reperfusion, as well as many technical factors. The formation, and subsequent modifications, of the Stroke Therapy Academic Industry Roundtable (STAIR) guidelines aim to increase translation of stroke therapeutics to the patient population by correcting for many of the technical limitations of prior studies ^14,15^. Thus, there is a great need for developing additional therapeutics for the stroke population that are targeted to the downstream molecular pathways that lead to tissue loss after ischemia and evaluating these new therapeutic candidates using the STAIR guidelines. These new agents may serve as adjuvants to currently available revascularization therapies. Additionally, because these pathways are active in the days following stroke, new agents targeted against neuroinflammation and cell death may have longer therapeutic administration windows than current revascularization therapies, may have fewer exclusion criteria or contraindications than the current therapies, or may limit the secondary side effects of stroke that are not treatable by the current revascularization therapies alone.

Central to the immune response is the nuclear factor kappa B (NF-κB) cascade. NF-κB is a transcription factor that regulates the expression of many pro-inflammatory and pro-apoptotic genes. In its inactive form, NF-κB is restricted to the cytoplasm of cells and is bound by its inhibitor, the inhibitor of kappa B (I-κB). However, upon activation by many diverse stimuli like IL-1, IL-6, TNF-α, and the activity of toll-like receptors (TLRs), I-κB becomes phosphorylated and ubiquitinated for proteasomal degradation, allowing the active subunits of NF-κB to enter the nucleus of the cell and enhance transcription of downstream pro-inflammatory and pro-apoptotic genes ^16–18^. Therefore, inhibition of NF-κB may be therapeutically beneficial by reducing the secondary tissue injury that occurs due to inflammation following ischemic stroke. A large number of preclinical studies have demonstrated the benefit of NF-κB inhibition on stroke outcome, but these results have not yet translated to the clinic. Trials using anti-inflammatory agents like minocycline and natalizumab have not shown benefit in the patient population, but specific inhibition at the level of the master regulator of inflammation and apoptosis, NF-κB, and timely intervention may result in improvements long-term in the stroke population ^14,15^.

Our lab has developed inhibitors of the NF-κB cascade attached to a protein drug carrier elastin-like polypeptide (ELP). ELPs consist of a series of amino acid repeats taken from human tropoelastin, are non-immunogenic ^19^ and thermally responsive ^20^, and stabilize fused peptide therapeutics in the systemic circulation ^21,22^. ELPs can also be modified to enhance delivery and target specific pathways for treatment of various diseases, including breast cancer ^23^, glioma ^24–27^, preeclampsia ^22^, and renal disease ^28^. For the current study, the N-terminus of ELP was modified with a cell-penetrating peptide, SynB1^29^, which is non-toxic and aids in the transport of the protein across the plasma membrane to the cells’ cytoplasm ^30^. The ELP domain consists of a series of VPGxG repeats that can be customized to alter the size and transition temperature of the protein ^31^. The C-terminus of ELP was modified with specific peptide inhibitors of the NF-κB cascade ^21,32–36^. The goal of this study was to create several inhibitors of the NF-κB cascade using ELP-peptide fusion technology, determine the optimal ELP-fused peptide for NF-κB inhibition, and test the agent for use as an adjuvant to revascularization for treatment of ischemic stroke.

## Methods

### Cloning, Purification, Labeling, and Characterization of SynB1-ELP – fused NF-κB Inhibitory Peptides

A pET25 vector encoding the cell penetrating peptide SynB1 underwent restriction digest using BamHI (Promega, catalog #R6021) for 1 hour at 37 °C and SfiI (New England Biolabs, catalog #R0123) for 5 hours at 50 °C. DNA cassettes for NF-κB inhibitory peptides p65P1, p65P6, and NEMO-leucine zipper were custom-designed (Integrated DNA Technologies) and were annealed by decreasing temperature from 95 °C to 20 °C at a rate of 0.5 °C/minute in an Eppendorf MasterCycler Gradient (catalog #5331). Subsequently, the constructs were ligated to the pET25 vector in separate reactions using T4 DNA Ligase (New England Biolabs, catalog #M0202) overnight at room temperature, transformed into XL1 Blue competent cells (Fisher, catalog #50-125-058) and plated on agar plates (2% LB Broth, Fisher, catalog #12780052) containing ampicillin (1%, Fisher, catalog #11593027). Colonies were selected and grown in liquid broth (Fisher, catalog #12780052) overnight at 37 °C for DNA isolation (by QIAprep Spin Miniprep Kit, Qiagen, catalog #27106) and sequence verification (EuroFins Genomics).

Next, an SfiI digest was conducted, and the ELP sequence was ligated to the vector (as mentioned above). The product was plated on an agar dish, and colonies were randomly selected, plasmid DNA was isolated, and the coding region was sequenced for verification to contain both the construct of interest and ELP sequence. In later experiments, SynB1-ELP was used to control for interactions that may be due to the cell penetrating peptide or ELP itself. This was created as mentioned above for the other constructs, but no NF-κB interacting element was attached ^21,32^. Once the DNA sequences were confirmed, they were transformed into BLR cells for protein expression using inverse transition cycling as described previously ^20,37^. For biodistribution and pharmacokinetics studies, proteins were fluorescently labeled on a unique cysteine residue using tetramethylrhodamine-5-maleimide as previously described ^38^.

### Cell Culture

RAW 264.7 cells (ATCC) were cultured in Dulbecco’s Modification of Eagle’s Medium (DMEM, Corning) and 10% fetal bovine serum and antibiotic/antimycotic. SH-SY5Y neurons (ATCC) were cultured in a 1:1 mixture of Eagle’s Minimum Essential Medium (EMEM, Lonza BioWhittaker) and F12 Medium (Gibco) with 10% fetal bovine serum and antibiotic/antimycotic. BV-2 microglia (Accegen) were cultured in Dulbecco’s Modification of Eagle’s Medium (DMEM, Corning) and 10% fetal bovine serum and antibiotic/antimycotic. For all cell types, experiments were conducted before cells reached 10 passages.

### Assessment of *In Vitro* Efficacy of ELP-NF-κB Inhibitory Peptides

RAW 264.7 murine macrophages were seeded in a 96-well plate at 15,000 cells/well. 24 hours later, media was removed and replaced with media containing with 1-50 µM of ELP-NF-κB inhibitory peptides (SynB1-ELP-p50i, SynB1-ELP-p65P1, SynB1-ELP-p65P6, or SynB1-ELP-NLZ, n = 3/protein concentration). The next day, media was removed, and the cells were rinsed with PBS. Media containing 10 µg/mL of lipopolysaccharide was applied to the cells for 1 hour before being removed and replaced with fresh media. The next day, media was removed, and secreted TNFα was measured by ELISA (Fisher, catalog #BMS607-3) using a Tecan plate reader at 450 nm. The remaining cells were incubated with media containing MTS reagent (Promega) for two hours before measuring absorbance at 490 nm using a Tecan plate reader.

### Assessment of SynB1-ELP-p50i Intracellular Localization

SH-SY5Y neurons and BV-2 microglia were seeded in 8-well chamber slides at 150,000 cells/well. The next day, cells were treated with 50 µM of rhodamine-labeled SynB1-ELP-p50i for 24 hours at 37 °C in a humidified incubator at 5% CO_2_. After treatment, cells were rinsed with PBS, fixed with 4% paraformaldehyde, and stained with DAPI (Fisher, catalog #62248). Intracellular localization was determined by imaging with a laser scanning confocal microscope (Nikon) with a 60x oil immersion objective using 405 nm and 561 nm lasers to excite DAPI and rhodamine, respectively. Images were collected using NIS-Elements imaging software and adjusted for brightness and contrast using Image J software. Images show representative intracellular distribution at a focal plane through the cell center.

### Animal Use

All protocols were approved by the University of Mississippi Medical Center Institutional Animal Care and Use Committee and in agreement with the National Institutes of Health Guidelines for Care and Use of Laboratory Animals. Male spontaneously hypertensive rats (SHR, Charles River) or Wistar-Kyoto rats (WKY, Charles River) were received at 11 weeks of age and allowed to acclimate for one week prior to use in experiments. Animals were maintained on a 12:12 hour light-dark cycle, at constant 23 °C temperature, and given food and water *ad libitum*.

### Middle Cerebral Artery Occlusion Surgery

Rats were anesthetized using isoflurane and temperature was maintained at 37 °C using homeothermic blankets. The common carotid, internal carotid, and external carotid arteries were isolated. The occipital artery and superior thyroid artery branches of the external carotid artery were cauterized and cut to allow for manipulation of the external carotid artery. After tying a distal suture around the external carotid artery, a filament (0.39 mm diameter, Doccol Corporation) was placed in the external carotid artery and advanced into the internal carotid artery until significant resistance was felt following the Longa method of intraluminal filament occlusion ^39,40^. Occlusion of the middle cerebral artery was verified by using a Laser Doppler Blood FlowMeter (AD Instruments) mounted to the skull at the MCA territory, bregma -1 mm and 5 mm lateral from the midline. The filament was tied in place for 2 hours of occlusion before removal for reperfusion. When the filament was removed, the external carotid artery was permanently tied, and the incision was sutured closed. Body temperature was maintained throughout the surgery at 37 °C using a homeothermic blanket with rectal probe. Sham surgeries consisted of all of the above without filament insertion.

### Mean Arterial Pressure (mmHg) Measurement

Prior to filament insertion, a catheter (V1 and V3 tubing, Scientific Commodities) was placed in the external carotid artery and advanced to the aortic arch. Mean arterial pressures (MAPs) were recorded using Cobe III pressure transducers (CDX Sema), and data were collected using receivers, amplifiers, and Lab Chart 7 PowerLab software from AD Instruments. MAPs were determined as the average of 1 minute of blood pressure recordings. The catheter was removed and discarded prior to filament insertion.

### Biodistribution and Pharmacokinetics of SynB1-ELP-p50i in SHRs

Following 2 hours of MCAO, SHRs remained anesthetized and were injected with rhodamine-labeled SynB1-ELP-p50i (50 mg/kg) either intravenously (IV, femoral vein) or intraarterially (IA, catheter placed in external carotid artery). Blood was collected via the tail vein at regular intervals for 4 hours post-injection to calculate plasma concentration over time. Prior to sacrifice, rats were injected with 30 mg/kg fluorescein isothiocyantate (FITC)-dextran (500 kDa, Sigma catalog #FD500S) to label the cerebrovasculature. Animals were euthanized, and brain, heart, liver, lungs, kidneys, and spleens were harvested and imaged using the In Vivo Imaging System (IVIS, Perkin Elmer). Regions of interest were drawn around each organ and the ipsilateral and contralateral hemispheres of the brain using Living Image software (Perkin Elmer) to determine average radiant efficiencies for each organ. Autofluorescence values were determined from imaging organs from a saline injected animal and were subtracted from the average radiant efficiency of each organ.

Blood samples were centrifuged, and plasma was transferred to a clean 1.5 mL tube. Fluorescence of plasma samples was directly measured using a Take 3 Microvolume Plate (Agilent), measured using an Agilent Cytation 7 (Excitation: 535, Emission: 580, Gain: 100), and fit to a standard curve of rhodamine-labeled SynB1-ELP-p50i.

### Acute Effects of SynB1-ELP-p50i on Infarct Volume and Gene Expression

Male SHRs were randomly assigned to three groups and received either sham or MCAO followed by treatment with salinen or 50 mg/kg SynB1-ELP-p50i. Sample size was determined to be n = 13/group using power analyses conducted with G*Power (RRID: SCR_013726). Animals were excluded if the reduction in cerebral blood flow was < 60 %.

#### Infarct Size Measurements

For a separate cohort of SHRs, following sham surgery or 2 hours MCAO, rats were treated with 50 mg/kg SynB1-ELP-p50i or saline IV (femoral vein, n = 13/group). 24 hours following reperfusion, blood was taken from the abdominal aorta, kidneys were harvested and fixed in 10% neutral buffered formalin (Fisher), spleens were harvested and weighed, and brains were harvested and sectioned into 2 mm sections using a brain slicing matrix. Brain sections were stained using 2% 2,3,5-triphenyltetrazolium chloride (TTC) at 37 °C for 15 minutes for visualization of the infarct. Two independent, experimentally blinded observers measured infarct sizes using Image J software by drawing regions of interest around the infarcted region on seven slices of each brain.

#### Gene Expression Analysis

RNA was isolated using TRIzol (Ambion). TTC-stained brain slices were sectioned into hemispheres and placed into Lysing Matrix D tubes (MP Biomedicals; Solon, OH) with 1 mL of TRIzol. The samples were homogenized with three 60 seconds bursts with 60 second breaks between each burst using a FastPrep-24 (MP Biomedicals). The liquid was transferred to new 1.5 mL centrifuge tubes with chloroform before centrifugation. The upper aqueous layer was transferred to a new 1.5 mL centrifuge tube with isopropanol. The tubes were centrifuged, aspirated, and washed with ethanol. The samples were air dried and dissolved in 30 µL of RNAse free water, concentrations were measured using a NanoDrop 2000 Spectrophotometer (Thermo Scientific), and RNA quality was assessed by capillary electrophoresis using an Agilent Bioanalyzer. RNA was converted to cDNA using the RevertAid First Strand cDNA Synthesis Kit (Thermo Fisher Scientific catalog #K1622) and instructions provided within the kit. A master mix of buffer, dNTP, and reverse transcriptase was made and added to each sample with oligo dT and RNA to make cDNA. The samples were incubated at 42 °C for 60 minutes before terminating the reaction by heating to 70 °C for 5 minutes. cDNA concentrations were measured using the NanoDrop 2000 Spectrophotometer (Thermo Scientific). Quantitative real-time PCR was performed on each hemisphere of TTC-stained brains from the infarct size comparison study by using Platinum II *Taq* Hot-Start DNA Polymerase (Thermo Fisher Scientific catalog #14966025) and its included protocol. PCR was performed using a C1000 Touch Thermal Cycler with the CFX96 Real-Time system head. Quantification was performed using the ΔΔCt method. All samples were measured in duplicate with β-actin loading controls to establish ΔCt values.

#### Measurement of Plasma Markers

Plasma from blood collected from the abdominal aorta was sent to the University of Mississippi Medical Center’s Analytical and Assay Core for analysis of markers of organ damage and toxicology. Alanine aminotransferase (ALT), aspartate aminotransferase (AST), blood urea nitrogen (BUN), creatinine (CREAT), and lactate dehydrogenase (LDH) levels were measured from the plasma samples using a VET AXCEL chemistry analyzer.

### Long-Term Effects of SynB1-ELP-p50i on Motor Recovery and Survival

Male SHRs were randomly assigned to four groups and received either sham or MCAO followed by treatment with saline, a single injection of SynB1-ELP-p50i (50 mg/kg), or daily injections of SynB1-ELP-p50i (50 mg/kg/day). Sample size was determined to be n = 15/group using power analyses conducted with G*Power (RRID: SCR_013726). Animals were excluded if the reduction in cerebral blood flow was < 60 % or if they died during the first 48 hours due to hemorrhagic transformation.

#### Behavioral Testing

A number of behavior tests and batteries were utilized to determine functional deficit and recovery in animals following MCAO or sham surgeries and treatment with either SynB1-ELP-p50i or saline. Animals were trained for three days prior to surgery, and behavioral deficits were assessed on days 1, 2, 3, 7, and 14 following stroke or sham surgery. Space and equipment were provided by the University of Mississippi Medical Center’s Animal Behavior Core.

##### Bederson Score

A modified Bederson Score was utilized to determine neurological deficit on days 1, 2, and 3 following MCAO or sham surgeries. Experimentally blinded observers scored behavior based on forelimb flexion, resistance to lateral push, and circling behavior as described previously ^41^. Scores ranged from 0, having no deficits, to 5, experiencing extreme neurological deficit.

##### Rotarod

Animals were trained for three days prior to surgery for performance on the rotarod. Training sessions included three trials of 60 seconds at a constant speed of 5 rpm. Animals were only included if they could stay upright and walking on the rod for all 60 seconds of a training trial. Testing occurred on days 1, 2, 3, 7, and 14 after MCAO and used an accelerating rod starting at 4 rpm and accelerating to 20 rpm over 300 seconds. Latency to fall, speed at fall, and distance traveled were recorded for each animal. The average of the three testing trials for each animal on each day was utilized in the statistical analysis.

##### Adhesive Removal Test

Animals were trained for three days prior to surgery for performance on the adhesive removal test. Briefly, paper dot stickers (0.64 cm diameter) were placed on the two forepaws of each animal. Time to remove the sticker from the left (ipsilateral) and right (contralateral) paw was recorded. Training and testing consisted of three trials daily for three days, and animals were included in the study if they could remove the stickers from their forepaws within 30 seconds by the end of the training period. For the testing period, animals had up to 180 seconds to remove the stickers from the forepaws.

##### Open Field Test

An Opto-Varimex Auto-Track 3 System (Columbus Instruments) was used to record open field behavior of the rats. Briefly, rats were placed in a 43 × 43 cm open field for 30 minutes on days 1, 2, 3, 7, and 14 post-MCAO. The system recorded total distance traveled, ambulatory time, resting time, and stereotypic behavior time. Data collected in the first 10 minutes of the open field test were used for analysis due to habituation to the environment.

#### Immunohistochemistry

On Day 14, animals were transcardially perfused using ice cold phosphate buffered saline and 4% paraformaldehyde (pH 7.4). Brains were harvested and soaked in 4% paraformaldehyde for 24 hours before being transferred to 30% sucrose prior to sectioning in 30 µm slices using a cryostat (Epredia CryoStar NX50). The primary antibodies were rabbit polyclonal ionized calcium-binding adapter molecule 1 (Iba1, 1:1500, FujiFilm Wako Pure Chemical Corporation catalog #019-19741) and mouse monoclonal RBFOX3/NeuN (1:1000, Novus Biologicals catalog #NBP1-92693). Secondary antibodies used were goat-anti-rabbit Alexa Fluor 488 (1:500, Thermo Fisher Scientific, catalog #A11070) and goat-anti-mouse Alexa Fluor 568 (1:500, Thermo Fisher Scientific, catalog #A11004). Sections were imaged using laser scanning confocal microscopy (Nikon C2+) using 405-, 488-, and 561-nm lasers for excitation of DAPI, Alexa 488, and Alexa 568, respectively. Images were taken in the penumbral region of the ipsilateral cortex to the MCAO (n = 3 images/animal) before being counted using ImageJ.

### Statistical Analysis

Results are expressed as mean ± SD. All statistical analyses were performed using GraphPad Prism 9 software, and a *p*-value < 0.05 was considered statistically significant. When appropriate, student’s t-test, one-way ANOVA, two-way ANOVA, and mixed-model analyses were utilized. Normality was assessed using a Shapiro-Wilk test prior to analysis. Survival after MCAO was analyzed using a Kaplan-Meier curve with a log-rank test.

## Results

### *In Vitro* Assessment of ELP-fused NF-κB Inhibitory Peptide Activity and Subcellular Localization

Four inhibitors of the NF-κB inflammatory cascade were screened for potency of NF-κB inhibition. The ELP-fused inhibitory peptides were designed to inhibit the trimerization of NF-κB Essential Modulator (NEMO) by mimicking the NEMO leucine zipper domain (NLZ)^35^, the phosphorylation of the p65 subunit of the NF-κB heterodimer (p65P1 and p65P6)^34^, or the nuclear localization of the activated NF-κB transcription factor by mimicking the p50 nuclear localization sequence (p50i)^21,22,36^ (Figure 1A). To assess NF-κB inhibition, the dose-response of each fusion protein’s ability to block LPS-stimulated TNF-α production in murine macrophages was assessed. SynB1-ELP-p50i was the most potent of the fusion proteins tested, showing significant reduction of TNF-α production at concentrations from 1 µM to 50 µM (Figure 1B) while causing no inhibition of cell proliferation (Figure 1C). This result, combined with previous data from our lab showing efficacy of SynB1-ELP-p50i to reduce NF-κB activation in HUVECs and RAW 264.7 cells^22^ led us to select SynB1-ELP-p50i as our lead candidate for further *in vitro* and *in vivo* assessment. The intracellular localization of SynB1-ELP-p50i was determined in SH-SY5Y neurons and BV-2 microglia. When cells were treated with 50 µM of rhodamine-labeled SynB1-ELP-p50i for 24 hours, SynB1-ELP-p50i localized to the cytoplasm of both cell types, indicating that the protein could cross the plasma membrane of both neurons and microglia and be readily available for NF-κB inhibition (Figure 1D and 1E).

**Figure 1:**
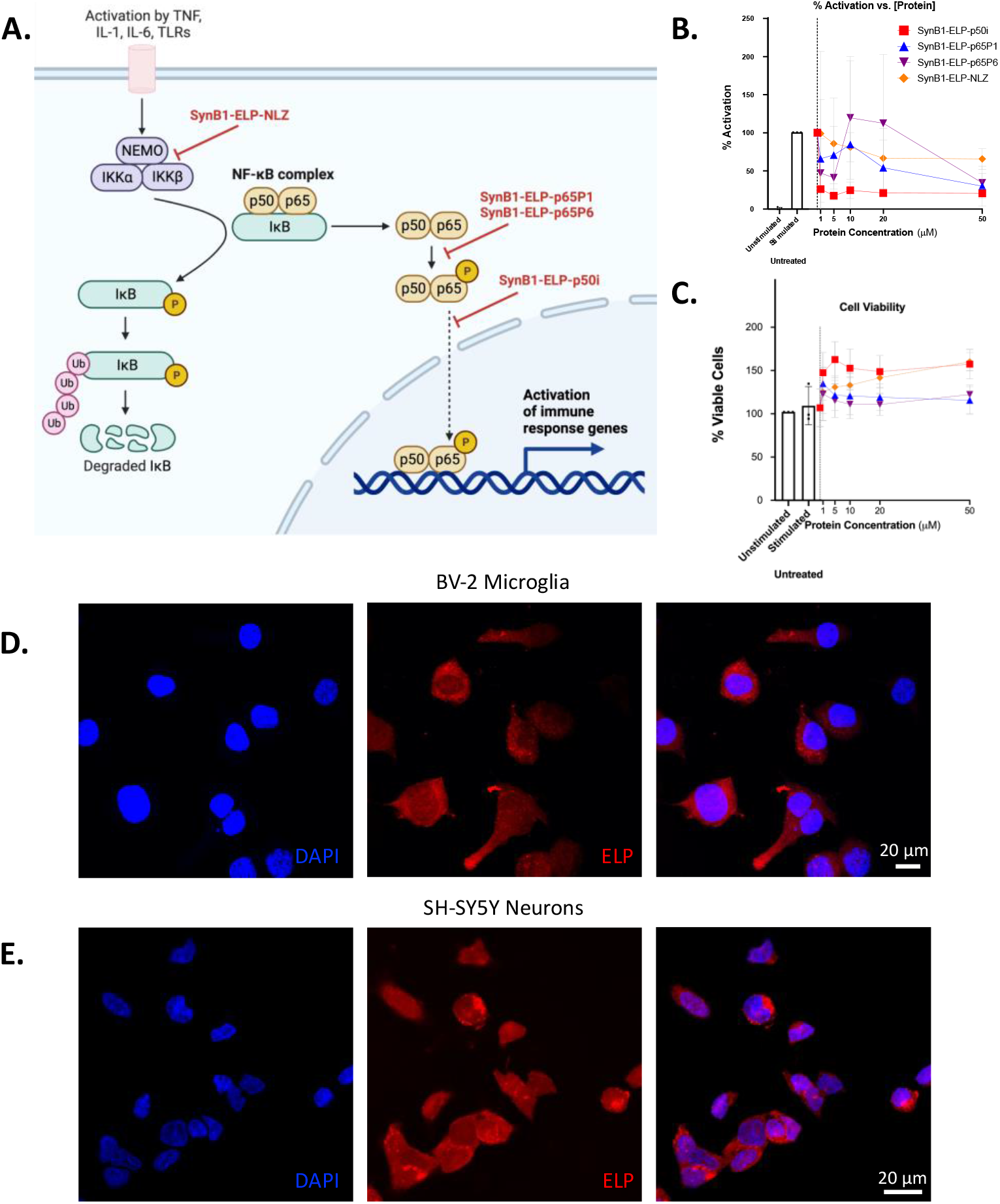
SynB1-ELP-p50i inhibits NF-*κ*B activation and localizes to the cytoplasm of cells. **A**. Schematic of NF-*κ*B pathway showing sites of inhibition of various ELP-fused NF-*κ*B inhibitory peptides. Adapted from “NF-KB Signaling Pathway”, by BioRender.com (2023). Retrieved from https://app.biorender.com/biorender-templates. **B-C**. RAW 264.7 cells pre-treated with 0-50 µM of SynB1-ELP-NF-*κ*B inhibitory peptides then stimulated with LPS (10 µg/mL) to activate the NF-*κ*B cascade. SynB1-ELP-p50i significantly reduced production of TNF-α at all doses compared to the untreated, stimulated control group (n = 3/protein concentration, one-way ANOVA, *F*(5,12) = 1.175, Dunnett’s multiple comparisons test, *p* < 0.0001) with no effect on cell viability. **D-E**. BV-2 microglia or SH-SY5Y neurons seeded 30,000 cells/well and treated with 50 µM of rhodamine labeled SynB1-ELP-p50i for 24 hours before being fixed and stained with DAPI. Imaging with confocal microscopy at 60x magnification showed localization of SynB1-ELP-p50i within the cytoplasm of the cells.

### Pharmacokinetics and Biodistribution of SynB1-ELP-p50i Following Ischemic Stroke

The biodistribution and pharmacokinetics of SynB1-ELP-p50i were assessed following middle cerebral artery occlusion (MCAO). Two routes of administration were compared, systemic intravenous injection (IV, femoral vein) and direct intraarterial injection (IA, left internal carotid artery) to determine which route of delivery led to the greatest deposition of SynB1-ELP-p50i in the brain, specifically the ipsilateral hemisphere to the MCAO. The IA route was used to test the hypothesis that local delivery, which would be clinically feasible following a mechanical thrombectomy, would result in delivery of higher therapeutic levels to the infarct site.

There was significant accumulation of SynB1-ELP-p50i in the ipsilateral hemisphere to the MCAO, consistent with deposition in the infarct and ischemic penumbra, following both IV and IA routes of delivery (Figure 2A and C, relative fluorescence quantified in Figure 2B and D). This accumulation is likely due to the weakening of the blood-brain barrier (BBB) following ischemic stroke and MCAO ^3,42,43^. SynB1-ELP-p50i is able to take advantage of the BBB permeability to enter the brain and localizes to the site of injury (Figure 2C, relative fluorescence quantified in Figure 2D). Interestingly, no differences were seen in the amount of protein present in the infarct four hours after IV versus IA delivery, refuting the hypothesis that direct delivery would result in better infarct targeting. In addition to localization at the infarct site, SynB1-ELP-p50i was present at high levels in the kidneys, consistent with its clearance by renal filtration as observed in our previous studies^24,31,44^. For quantitative assessment of the concentration of SynB1-ELP-p50i in the infarct, 30 µm thick slices of brain and protein standards were analyzed using quantitative fluorescence histology^45^. Fluorescent histology analysis showed that the 50 mg/kg dose used resulted in infarct concentration of ∼50-60 µg/mL (∼1.0 – 1.5 µM) (Figure 2E, quantified in Figure 2F). These intra-infarct concentrations were consistent with concentrations at which SynB1-ELP-p50i was active in *in vitro* NF-κB inhibition assays. As with the whole organ imaging, no differences were observed between IV and IA routes of delivery for deposition of SynB1-ELP-p50i in the infarct. Therefore, the remaining studies utilized systemic delivery. Direct measurement of plasma fluorescence and fitting to a two-compartment pharmacokinetic model revealed that SynB1-ELP-p50i had a terminal half-life of approximately 90 minutes (Figure 2G). There were no significant differences in plasma concentration over time following either an IV or IA injection of SynB1-ELP-p50i (Figure 2G). These results were validated and remained consistent in normotensive Wistar-Kyoto rats subjected to MCAO (Supplementary Figure 1).

**Figure 2:**
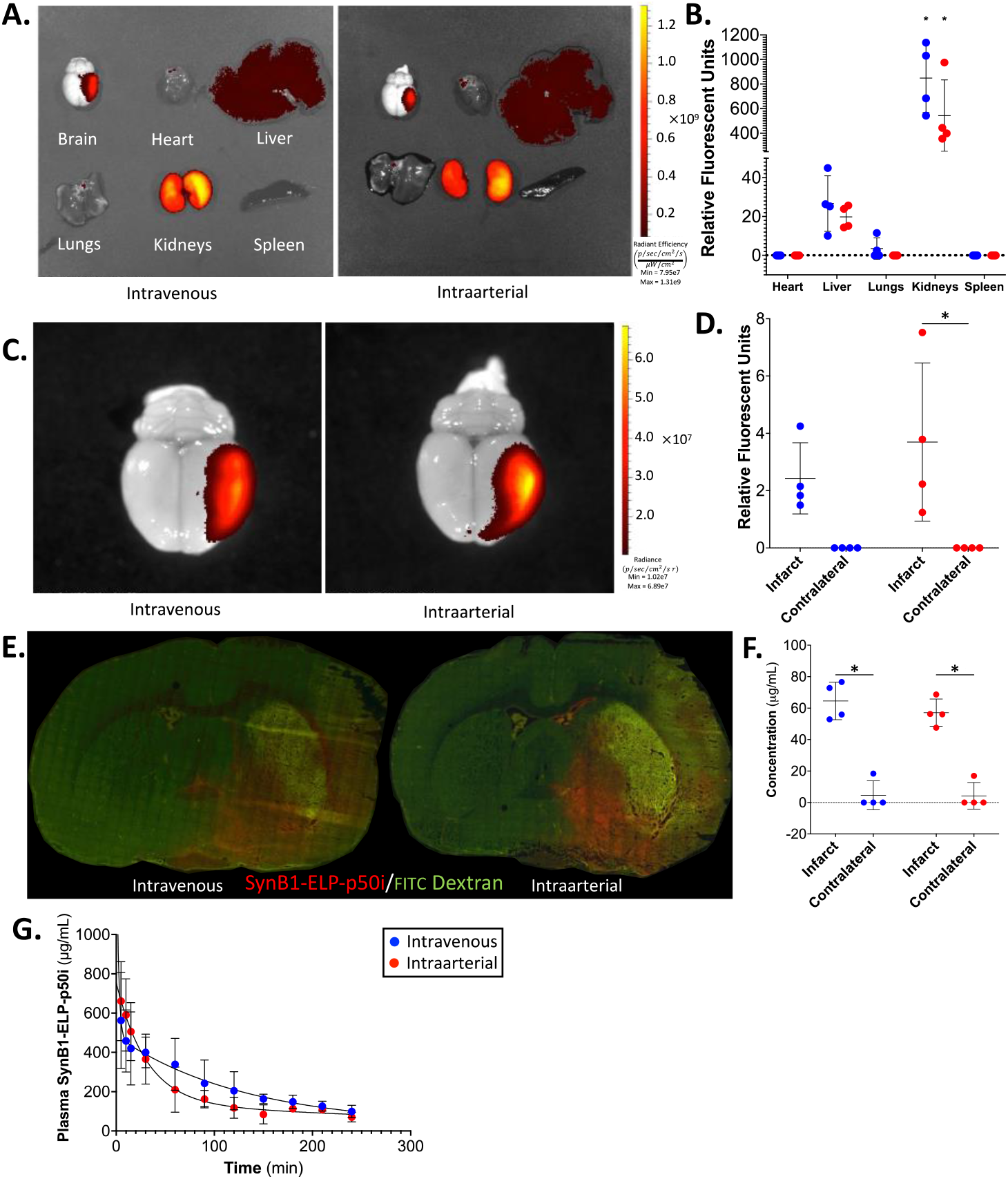
SynB1-ELP-p50i localizes to the infarct following intravenous or intraarterial injection after MCAO in SHRs. **A**. Representative IVIS images of IV and IA SynB1-ELP-p50i treated animals. **B**. Relative fluorescence of IV and IA SynB1-ELP-p50i treated animals shows high accumulation in the kidneys compared to every other organ (n = 4/route of delivery, two-way ANOVA; *F*(4,30) = 46.61, Tukey’s post-hoc, *p* < 0.0001). **C**. Representative images of brains of IV and IA SynB1-ELP-p50i treated animals in smaller field of view. **D**. Quantified relative fluorescence of brain deposition showed significant increase relative to contralateral brain of rhodamine labeled SynB1-ELP-p50i following IA delivery (two-way ANOVA; *F*(1,12) = 16.40, Šidák’s post-hoc, *p* = 0.0095). The difference between infarct and contralateral brain after IV delivery did not reach significance (*F*(1,12) = 16.40, *p* = 0.0827). **E**. Representative fluorescent images of 20 μm slices of brains following MCAO and treatment with 30 mg/kg FITC dextran to label vasculature (IV, femoral vein). **F**. Quantified fluorescent histology showing concentration of SynB1-ELP-p50i following IV or IA delivery. Both routes of delivery resulted in a significant increase in SynB1-ELP-p50i (two-way ANOVA; *F*(1,12) = 136.7, Šidák’s post-hoc, *p* < 0.001). **G**. Plasma concentration of rhodamine labeled SynB1-ELP-p50i following IV or IA injection showed no difference between the two routes of administration.

### Efficacy of SynB1-ELP-p50i to Reduce Infarct Size Following MCAO

After establishing that IV injection of 50 mg/kg SynB1-ELP-p50i resulted in potentially therapeutic levels of the therapeutic in the infarct, infarct volume was measured 24 hours post-MCAO in rats treated with SynB1-ELP-p50i versus saline treated MCAO and sham operated controls. There were no differences between Sham + Saline, MCAO + Saline, or MCAO + SynB1-ELP-p50i groups in pre-stroke body weight (Figure 3A) or mean arterial pressure (Figure 3B). Filament insertion depth (Figure 3C) and reduction in cerebral blood flow (Figure 3D) were not different between MCAO + Saline and MCAO + SynB1-ELP-p50i groups. MCAO animals treated with saline had large cortical infarcts averaging 373.74 ± 48.15 mm^3^. Infarct volume was significantly reduced in MCAO rats treated with SynB1-ELP-p50i (Figure 3E, quantified in Figure 3F). The 329.43 ± 60.21 mm^3^ infarct volume in SynB1-ELP-p50i treated rats represented a reduction in infarct volume of 11.86 %. Twenty four hours after stroke and treatment, the animals that were subjected to MCAO had a significant decrease in spleen weight compared to Sham + Saline animals, a phenomenon attributed to rehoming of resident immune cells from the spleen to the site of injury in the brain ^46,47^.There were no differences between MCAO + Saline and MCAO + SynB1-ELP-p50i in spleen weight (Supplementary Figure 2C and Supplementary Figure 2D).

**Figure 3:**
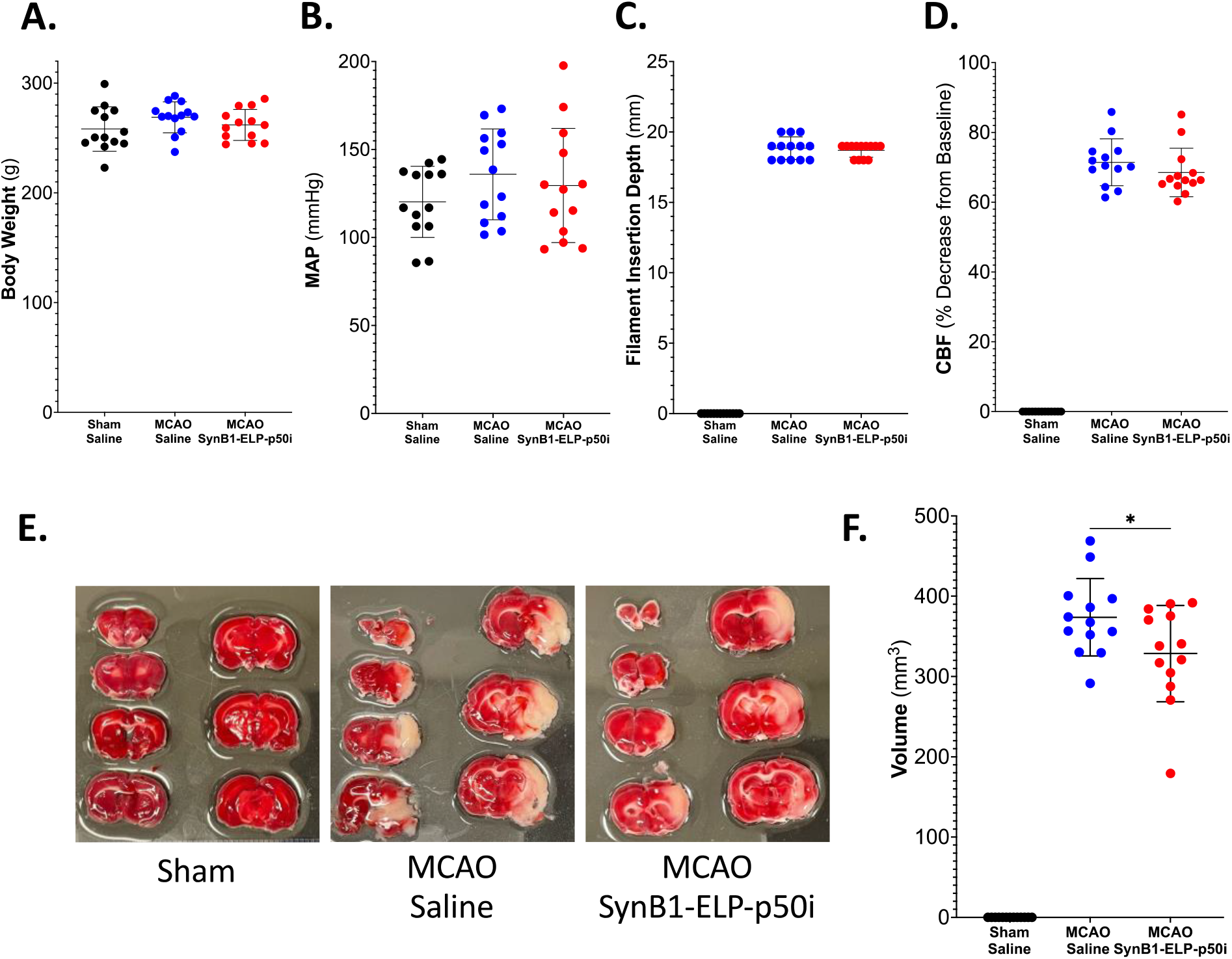
SynB1-ELP-p50i treatment reduces infarct size 24 hours following MCAO. **A**. There were no differences among groups in pretreatment body weight (one-way ANOVA; *F*(2,36) = 1.425, *p* = 0.2536). **B**. There were no differences among groups in pretreatment MAP (one-way ANOVA; *F*(2,36) = 1.143, *p* = 0.3301). **C**. There was no difference in insertion depth of the filament into the internal carotid artery between two MCAO groups (two-tailed, unpaired t test; *t*(24) = 0.5941, *p* = 0.5580). **D**. There was no difference in percent decrease in cerebral blood flow between MCAO groups measured using laser doppler flowmetry (two-tailed, unpaired t test; *t*(24) = 1.080, *p* = 0.2910). **E**. Representative images of TTC stained brains showed metabolically active tissue in red and metabolically inactive (infarcted tissue) in white. There was no observable infarct in the Sham + Saline group, but there was obvious infarction in both of the MCAO groups, with SynB1-ELP-p50i treated rats having smaller infarcts than saline treated rats. **F**. Treatment with SynB1-ELP-p50i significantly reduced infarct size compared to the MCAO + Saline treated animals (two-tailed, unpaired t test; *t*(24) = 2.073, *p* = 0.0491).

At the time of sacrifice, blood was collected from the abdominal aorta and used for measures of toxicology and organ dysfunction. There were no differences among groups in levels of plasma markers for liver damage, alanine aminotransferase or aspartate aminotransferase; kidney damage, blood urea nitrogen or creatinine; or general organ dysfunction, lactate dehydrogenase (Table 1). Despite high accumulation of SynB1-ELP-p50i in the kidneys, there were no histological signs of renal injury and no kidney fibrosis (Supplementary Figure 2A and 2B). Additionally, brains were sectioned into hemispheres and homogenized for qPCR analysis of genes affected by MCAO or downstream of NF-κB. MCAO induced a significant rise in IL-1β, IL-6, ICAM-1, CXCL-1, CXCL-2, and MMP-9. There were no statistically significant differences between saline and SynB1-ELP-p50i treated MCAO groups in gene expression, though there were trends for reduction of some pro-inflammatory genes following SynB1-ELP-p50i treatment (Supplementary Figure 3).

**Table 1:** SynB1-ELP-p50i treatment does not affect levels of organ damage markers 24 hours post-MCAO. Levels of markers for liver damage (ALT, AST), kidney damage (BUN, CREAT), and organ dysfunction (LDH) were measured 24 hours post-MCAO and showed no differences among groups, with the exception of CREAT. Plasma creatinine was significantly elevated in the MCAO SynB1-ELP-p50i treated animals compared to shams (one-way ANOVA; *F*(2,36) = 4.730, Tukey’s post-hoc, *p* < 0.05); however, the elevated value (0.35 ± 0.06) was still within normal physiological range.

|  | Sham | MCAO |  |
| --- | --- | --- | --- |
|  | Saline | Saline | SynB1-ELP-p50i |
| ALT, U/L | 32.62 ± 21.58 | 22.31 ± 15.31 | 30.23 ± 14.54 |
| AST, U/L | 208.08 ± 44.90 | 174.62 ± 98.59 | 231.23 ± 79.40 |
| BUN, mg/dL | 17.54 ± 3.36 | 21.46 ± 3.84 | 20.00 ± 5.73 |
| CREAT, mg/dL | 0.27 ± 0.08 | 0.34 ± 0.07 | 0.35 ± 0.06 * |
| LDH, U/L | 1632.69 ± 962.33 | 760.31 ± 830.57 | 1662.69 ± 1025.91 |

### Effects of SynB1-ELP-p50i on Animal Survival, Neurological Deficits, and Behavioral Outcomes following MCAO

For long-term assessment of potential toxicity and neurological and behavioral benefits of SynB1-ELP-p50i treatment, a 14-day efficacy study was conducted. SHRs received either a sham or MCAO surgery followed by IP injection with saline, a single dose of SynB1-ELP-p50i, or daily injections of SynB1-ELP-p50i. IP delivery was utilized to reduce stress and surgical burden on the animals throughout the study. Equivalent delivery of SynB1-ELP-p50i to the infarct site following IP injection and an equal biodistribution profile following MCAO in SHRs was verified at 4- and 24-hours post-injection and is shown in Supplementary Figure 4. At the time of surgery, there were no differences between groups in mean arterial pressure (Figure 4A), filament insertion depth (Figure 4B), or reduction in cerebral blood flow (Figure 4C). All MCAO groups, regardless of treatment condition, lost significant body weight compared to the Sham + Saline controls, but MCAO groups regained their weight between days 8 and 10 post-MCAO (Figure 4D). Treatment with a single dose of SynB1-ELP-p50i immediately following reperfusion improved survival for 14 days following MCAO from 55.56 % in MCAO + Saline treated rats to 63.64 % in MCAO + SynB1-ELP-p50i treated rats. Interestingly, the survival benefit was not observed when SynB1-ELP-p50i treatment was continued daily for the remainder of the experimental period (Figure 4E).

**Figure 4:**
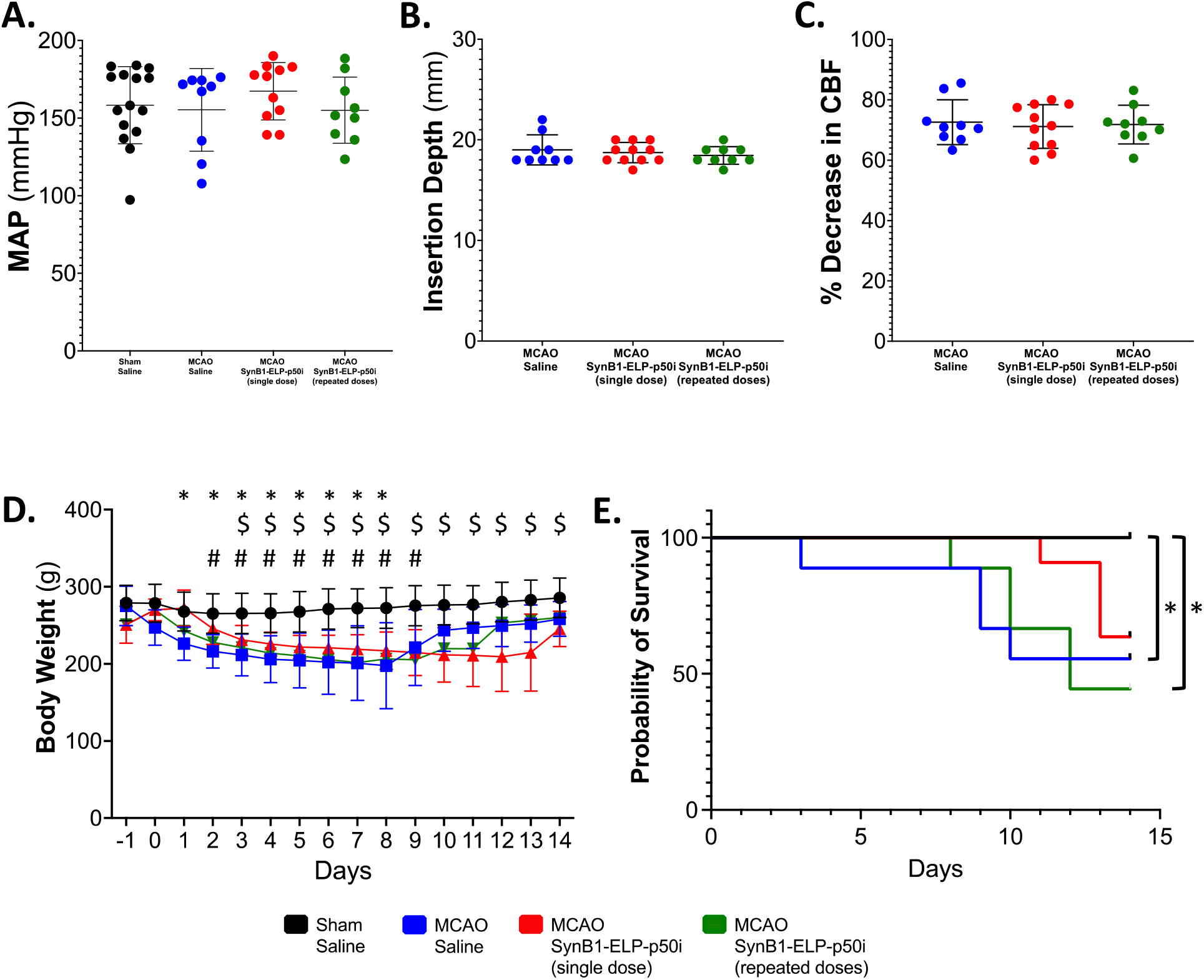
SynB1-ELP-p50i improves survival following MCAO. **A**. There were no differences between groups in pretreatment MAP (one-way ANOVA; *F*(3, 40) = 0.6380, *p* = 0.5950). **B**. There was no difference in insertion depth of the filament into the internal carotid artery between two MCAO groups (one-way ANOVA; *F*(2, 26) = 0.5249, *p* = 0.5978). **C**. There was no difference in percent decrease in cerebral blood flow between MCAO groups measured using laser doppler flowmetry (one-way ANOVA; *F*(2, 26) = 0.3309, *p* = 0.9022). **D**. MCAO + SynB1-ELP-p50i treatment did not significantly alter body weight compared to MCAO + Saline treated animals, but all MCAO animals had decreased body weight from day 3 to day 9 compared to the Sham + Saline treated animals (mixed-effects model (REML), Tukey’s multiple comparisons test; *F*(3, 41) = 12.67, Sham + Saline vs. * MCAO + Saline, $ MCAO + SynB1-ELP-p50i single dose, # SynB1-ELP-p50i repeated dose, *p* < 0.05). **E**. Kaplan-Meier survival analysis for animals following sham or MCAO surgery and treatment with saline or SynB1-ELP-p50i. Both the MCAO + Saline and MCAO + SynB1-ELP-p50i (repeated doses) group had significantly decreased survival compared to Sham + Saline controls (Log-rank Mantel-Cox with adjusted α_Bonferroni_ for multiple comparisons, * = p < 0.0083).

The Bederson score of neurological deficit was utilized on days 1, 2, and 3 to observe forelimb flexion, circling behavior, and resistance to lateral push^41^. While stroke caused significant deficits, there were no differences in MCAO groups regardless of treatment (Supplementary Figure 5A). Rotarod was used to measure changes in locomotor activity, and there were no differences in latency to fall between MCAO groups regardless of treatment condition (Supplementary Figure 5B). The adhesive removal test, used to measure sensorimotor deficits, showed impairment in all stroked rats but no differences among MCAO groups regardless of treatment condition (Supplementary Figure 5C). An open field test for measurement of general locomotor activity also showed no differences in any of the groups, with the exception of the MCAO + SynB1-ELP-p50i single dose group walking more distance on day 7 than the Sham + Saline controls (Supplementary Figure 5D).

On day 14, blood was collected from the heart and used for analysis of toxicology markers and markers of organ dysfunction. There were no differences among groups in levels of plasma markers of liver damage, alanine aminotransferase and aspartate aminotransferase (Table 2 and Supplementary Figure 6A and 6B). There were also no differences in blood urea nitrogen or plasma creatinine, indicating no renal damage, or in lactate dehydrogenase, a marker of general organ dysfunction (Table 2 and Supplementary Figure 6C-E). Also on day 14, the kidneys were harvested for Masson’s trichrome staining to detect renal fibrosis, and the spleen was harvested and weighed to observe any differences in spleen weight as a proxy for peripheral immune response to the MCAO. There was no detectable renal fibrosis (Supplementary Figure 7A and 7B). There were also no differences in spleen weight (Supplementary Figure 7C) or spleen index (Supplementary Figure 7D).

**Table 2:** SynB1-ELP-p50i treatment does not affect levels of organ damage markers 14 days post-MCAO. Levels of markers for liver damage (ALT, AST), kidney damage (BUN, CREAT), and organ dysfunction (LDH) were measured 14 days post-MCAO and showed no differences among groups.

|  | Sham | MCAO |  |  |
| --- | --- | --- | --- | --- |
|  | Saline | Saline | SynB1-ELP-p50i Single Dose | SynB1-ELP-p50i Repeat Doses |
| ALT, U/L | 61.20 ± 18.55 | 51.13 ± 11.17 | 53.25 ± 10.00 | 59.80 ± 19.84 |
| AST, U/L | 193.67 ± 195.70 | 128.75 ± 43.15 | 205.13 ± 92.71 | 222.80 ± 111.55 |
| BUN, mg/dL | 22.00 ± 5.45 | 21.38 ± 2.00 | 23.50 ± 2.45 | 21.20 ± 2.39 |
| CREAT, mg/dL | 0.35 ± 0.12 | 0.37 ± 0.06 | 0.39 ± 0.09 | 0.33 ± 0.11 |
| LDH, U/L | 373.60 ± 315.08 | 329.13 ± 213.97 | 501.63 ± 196.79 | 481.60 ± 233.40 |

On day 14, brains were stained with markers for microglia (Iba1) and neurons (NeuN). There were trends for increased microglial and decreased neuron numbers in rats subjected to stroke, and trends for normalization of these parameters in rats treated with a single dose of SynB1-ELP-p50i immediately after reperfusion, though the values did not reach statistical significance. (Figure 5A, quantified in Figure 5B and 5C). Interestingly, microglia number was significantly elevated in rats that received daily SynB1-ELP-p50i doses compared to the single dose group and to sham operated animals.

**Figure 5:**
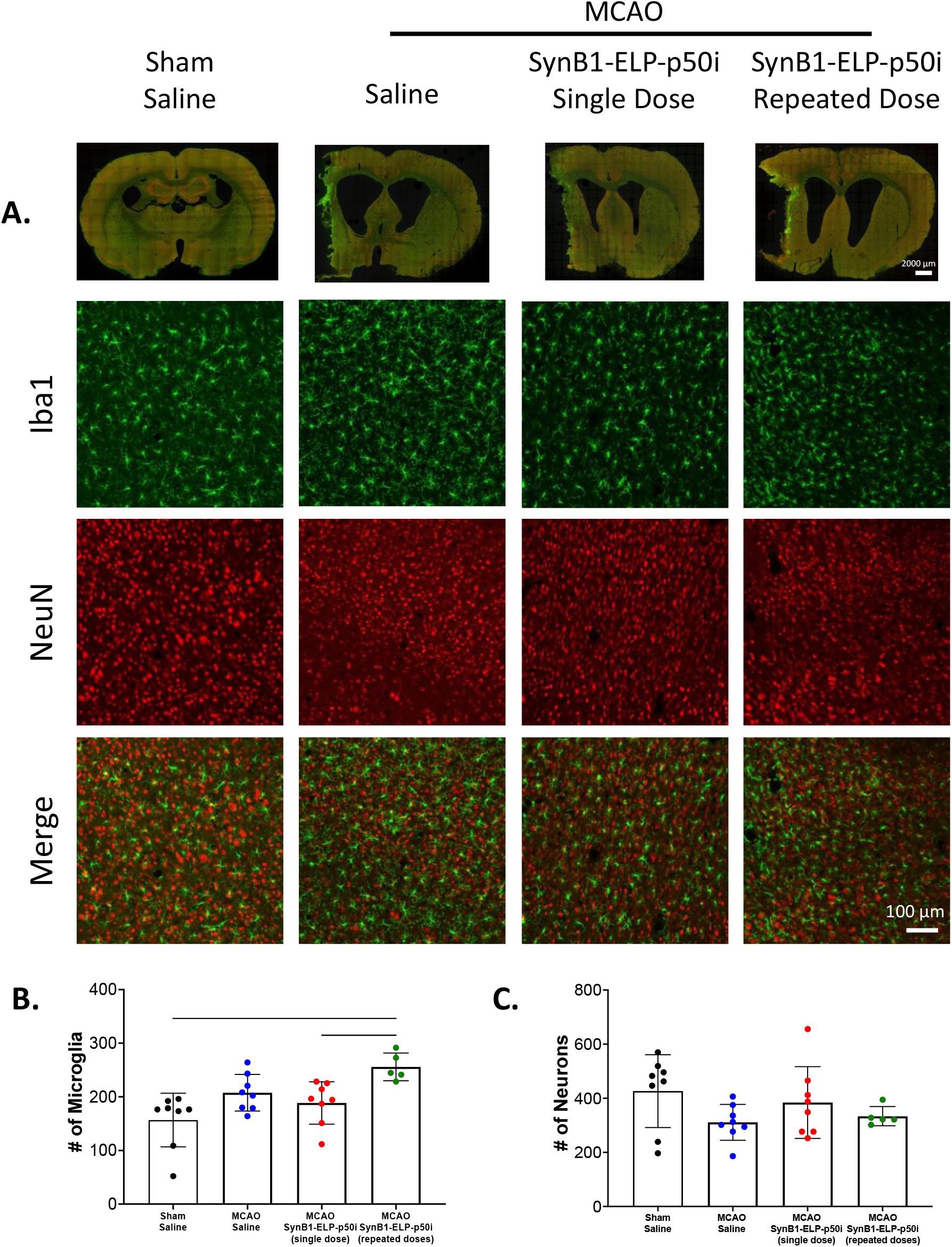
SynB1-ELP-p50i treatment does not affect Iba1 or NeuN positive cells. **A**. Whole slice stitches and 20x representative images of 30 μm sections stained with Iba1 and NeuN. **B**. Number of microglia in each group showed that SynB1-ELP-p50i repeated doses had the highest number of Iba1+ microglia, a significant increase compared to Sham + Saline and SynB1-ELP-p50i single dose (one-way ANOVA; *F*(3, 25) = 6.711, Tukey’s post-hoc, *p* < 0.05). **C**. Number of NeuN+ neurons showed no difference among groups.

## Discussion

The novel biologic SynB1-ELP-p50i is potent and effective at reducing NF-κB activation *in vitro* and is able to cross the plasma membrane and localize to the cytoplasm in neurons and microglia. Following both an IV and IA injection, SynB1-ELP-p50i is able to cross the BBB and localize to the infarct region following MCAO. As both of these routes are translationally relevant, whether a patient receives an IV thrombolytic or an endovascular thrombectomy, it is promising that SynB1-ELP-p50i can accumulate in the brain at therapeutically beneficial concentrations following either route of administration. Additionally, SynB1-ELP-p50i treatment significantly reduced infarct size 24 hours following MCAO, suggesting that the inhibition of NF-κB acutely by SynB1-ELP-p50i has therapeutic benefit.

In a 14-day longitudinal study, a single dose of SynB1-ELP-p50i improved survival compared to saline or animals given daily doses of SynB1-ELP-p50i. The differential benefit between the single dose and repeated dose paradigms demonstrates that the timing and duration of treatment is critical for an anti-inflammatory agent like SynB1-ELP-p50i. This is likely because inflammation following stroke has multiple phases, with antagonistic properties in the acute phase of stroke recovery but beneficial effects and repair in the chronic phase of stroke recovery^7,9,48,49^. Therefore, inhibiting inflammation in the early days after stroke is more beneficial than an extended continuous inhibition. While two treatment schedules were tested in this study, future experiments are needed to determine the optimal treatment window. It may be possible to induce further reductions in infarct size and better improvement in survival by treating for longer than one day or at varying doses, however the temporal threshold at which SynB1-ELP-p50i begins to negatively affect beneficial inflammation at more distant time points after stroke must be determined.

It is promising that, even after 14 days of continuous treatment and in spite of the very high renal deposition, SynB1-ELP-p50i did not cause any observable toxicity, as measured using both plasma markers of hepatic, renal, and general organ damage and by renal histology. Since SynB1-ELP-p50i is filtered by the kidneys and accumulates within the kidneys, it is important to be sure that there is no negative effect on kidney function or morphology as a result. By assessing plasma creatinine, blood urea nitrogen, and renal fibrosis, we were able to demonstrate that SynB1-ELP-p50i did not negatively affect the kidney when being used to treat inflammation following ischemic stroke.

Additionally, in the 14-day longitudinal study, there was little observable benefit of treatment with SynB1-ELP-p50i on functional and neurological recovery following 120 minutes of MCAO. There are many reasons why the efficacy of SynB1-ELP-p50i may not be seen in these long-term behavior studies. The most likely explanation is the occlusion time utilized in this study. One hundred and twenty minutes of occlusion leads to a large amount of ischemic insult in rats, especially in SHRs which have reduced collateral blood flow due to smaller collateral vessels with reduced vasodilatory capacity^50^. Additionally, SHRs are more susceptible to ischemic injury than normotensive rats, and MCAO in SHRs results in larger infarctions due to alterations that chronic hypertension induces on the cerebrovasculature and BBB as well as marked increases in oxidative stress and inflammatory signalling^51–53^. Future studies should be conducted using a shorter time of occlusion, 60-90 minutes, to determine if SynB1-ELP-p50i treatment can result in functional improvement in animals following a less severe stroke. While infarct size was reduced by SynB1-ELP-p50i treatment, it is possible that even these improved infarcts were still too severe to allow for observation of motor function recovery. It is possible that attrition in the saline-treated group due to rats succumbing to the stroke may have masked any benefits in motor function induced by SynB1-ELP-p50i treatment. It is also important to note that SynB1-ELP-p50i treatment did not worsen performance in behavior tasks.

While ELPs have been used preclinically in glioma^24–27^ models and intranasal administration^54^ for delivery to the central nervous system, to our knowledge, they have never been utilized as a drug carrier for delivery in models of ischemic stroke. The physical properties of ELPs are highly plastic, as their size, transition temperature, and hydrophobicity can be modified by changing the number of repeats or residue composition of the ELP sequence^31,55^. Our results showing high accumulation in the ischemic hemisphere of the brain are very promising for the future of ELPs as drug carriers for models of central nervous system disease, specifically those in which the BBB is weakened either acutely or chronically. By modifying the size of the ELP and the therapeutic peptide that is attached, the ELP drug delivery system may be used to treat other pathways implicated in diseases such as ischemic stroke.

## Data Availability Statement

The data that support the findings of this study are available on request from the corresponding author.

## Author Contributions

J.A. Howell and G.L. Bidwell conceived and designed the studies. J.A. Howell, E. Perkins, N. Gaouette, and M. Lopez performed the research and acquired the data. J.A. Howell, M. Lopez, and S. Burke analyzed and interpreted the data. All authors were involved in writing and revising the manuscript.

## Acknowledgements

We would like to acknowledge Drazen Raucher for SynB1-ELP and SynB1-ELP-p50i constructs. We would also like to thank Rowshan Begum for assisting in protein purification. *Ex vivo* fluorescence imaging of organs was performed using the University of Mississippi Medical Center Animal Imaging Core. Analysis of plasma concentrations of toxicology markers was conducted by the University of Mississippi Medical Center Analytical and Assay Core, which is supported by NIH grant P20GM104357. Kidney histology was conducted by the University of Mississippi Medical Center Histology Core, which is supported by NIH grant P20GM104357. Animal behavior testing was conducted using equipment provided by the University of Mississippi Medical Center Animal Behavior Core.

