## Supplemental Data for "Elastin-like polypeptide delivery of anti-inflammatory peptides to the brain following ischemic stroke"

**Biodistribution and Pharmacokinetics of SynB1-ELP-p50i in Normotensive Rats**

When normotensive female Wistar-Kyoto rats were treated with 50 mg/kg rhodamine-labeled SynB1-ELP-p50i, the protein accumulated at high levels in the brain, specifically in the infarcted region, following both an IV (femoral vein) and IA (internal carotid artery) injection. The protein also accumulated at high levels in the kidneys. There was no significant difference between routes of administration in organ deposition or pharmacokinetic clearance curves.

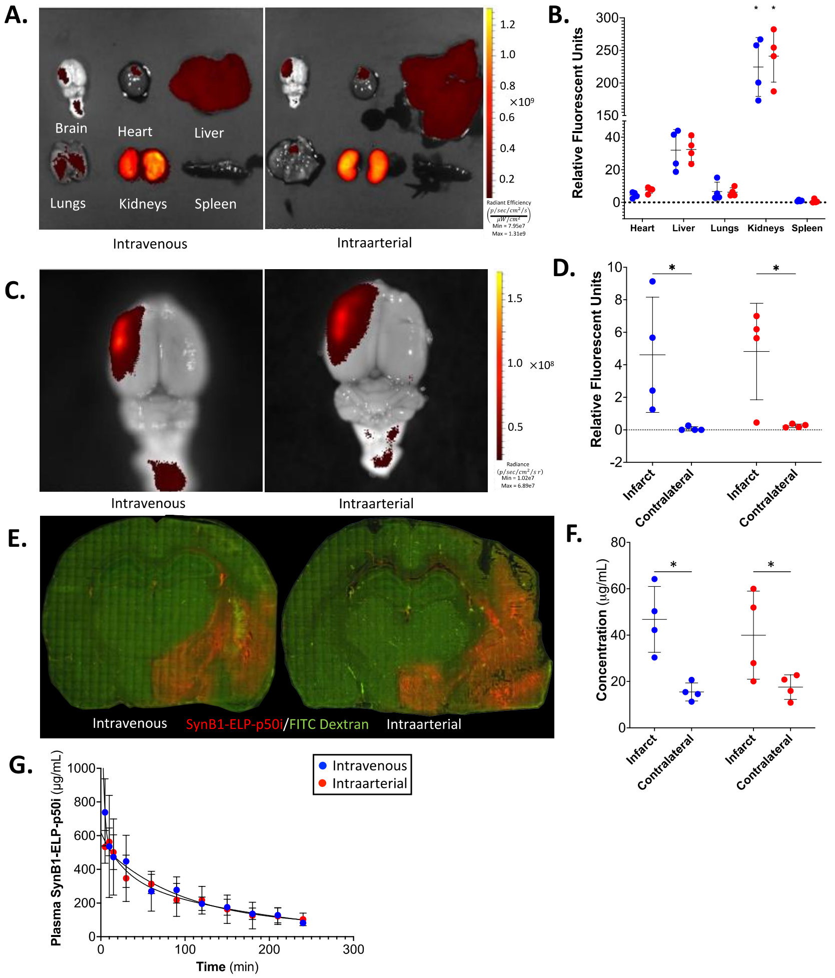


**Supplementary Figure 1: SynB1-ELP-p50i localizes to the infarct following intravenous or intraarterial injection after MCAO in Wistar-Kyoto rats. A.** Representative IVIS images of IV and IA treated animals. **B.** Quantified relative fluorescence of IV and IA treated animals showed high accumulation in the kidneys compared to every other organ (two-way ANOVA; $F(6,42) = 180.9$, $p < 0.0001$). **C.** Representative images of brains of IV and IA treated animals in smaller field of view. **D.** Quantified relative fluorescence of brain deposition showed significant increase of SynB1-ELP-p50i in the infarcted hemisphere following IV and IA delivery (two-way ANOVA; $F(1,12) = 15.58$, $p = 0.0019$). **E.** Representative fluorescent images of 20 µm thick slices of brains following MCAO and treatment with 30 mg/kg FITC dextran to label vasculature (IV, 30 mg/kg, femoral vein). **F.** Quantified fluorescent histology showing concentration of SynB1-ELP-p50i in the infarct and contralateral hemisphere IV or IA delivery resulted in a significantly higher deposition of SynB1-ELP-p50i (two-way ANOVA; $F(1,12) = 18.98$, Šidák's post-hoc, * $p < 0.05$). **G.** Plasma concentration of SynB1-ELP-p50i showed no difference between the two routes of administration.

**Acute Effects of SynB1-ELP-p50i on Renal Fibrosis and Spleen Weight**

Treatment with SynB1-ELP-p50i following MCAO did not cause fibrosis in the kidneys at 24 hours post-stroke, despite having a high intrarenal deposition. MCAO caused a decrease in spleen weight, due to resident splenocytes going to the site of injury in the brain. There were no differences between MCAO groups in spleen weight or spleen index, but both groups weighed significantly less than shams.

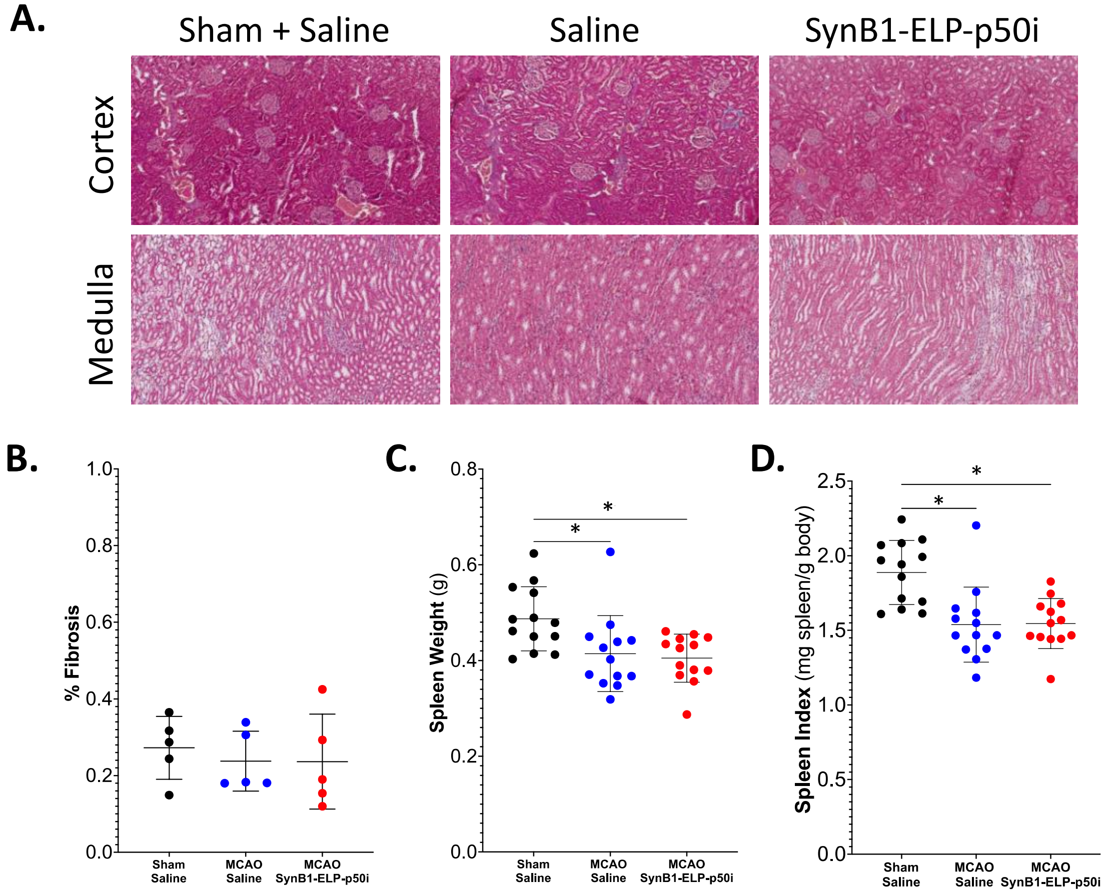


**Supplementary Figure 2: No kidney fibrosis or improvement in spleen weight seen 24 hours following MCAO and treatment with saline or SynB1-ELP-p50i. A.** Kidneys from 5 rats from each group were fixed, embedded in paraffin, sectioned using a microtome, and Masson's trichrome stained. Representative images are shown from the renal cortex and medulla for each group. Collagen is stained blue in these images and is representative of fibrosis that has occurred in the kidneys. Dark red shows cytoplasm or erythrocytes. **B.** There were no differences between groups in % Fibrosis for the entire kidneys (one-way ANOVA; $F(2,12) = 0.2217$, $p = 0.8044$). **C.** Spleen weight was measured using an analytical balance at time of sacrifice 24 hours following induction of MCAO and treatment with saline or SynB1-ELP-p50i (IV, 50 mg/kg). There were no differences between MCAO + Saline and MCAO + SynB1-ELP-p50i groups (one-way ANOVA; $F(2,36) = 5.930$, $p = 0.9311$), but both groups differed from Sham + Saline controls (Tukey's post-hoc, $p < 0.05$). **D.** To correct for any differences in body weight, spleen index was calculated by dividing the spleen weight (mg) by the animal body weight (g). There were no differences between MCAO + Saline and MCAO + SynB1-ELP-p50i groups (one-way ANOVA; $F(2,36) = 11.32$, Tukey's post-hoc, $p = 0.9950$), but both groups differed from Sham + Saline controls ($p < 0.05$).

**Gene Expression Changes Following Stroke and SynB1-ELP-p50i Treatment**

Expression of cytokines IL-1β, IL-6, CXCL1, and CXCL2 were increased in the ipsilateral hemisphere of MCAO animals. Treatment with 50 mg/kg SynB1-ELP-p50i did not affect gene expression of downstream targets of NF-κB as measured by qPCR, though there were trends for reduction of CXCL2 that did not reach statistical significance. Matrix metalloproteinase 9 (MMP-9), an enzyme involved in the blood-brain barrier breakdown, and intracellular adhesion molecule-1 (ICAM-1), a molecule upregulated post-stroke that leads to leukocyte accumulation, were also increased in the ipsilateral hemisphere post-stroke. There were also trends for reduced expression of these factors in rats treated with SynB1-ELP-p50i, but the reductions did not reach statistical significance.

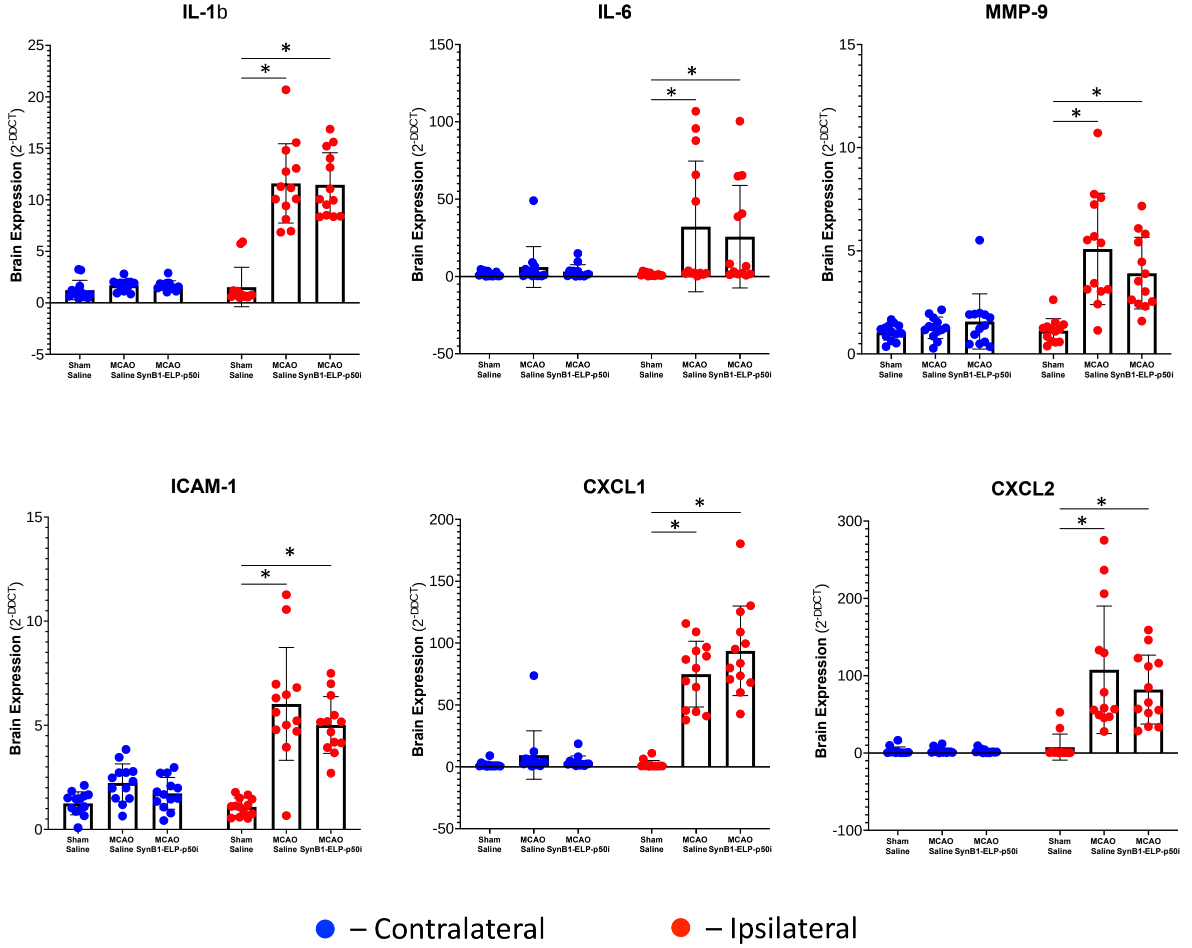


**Supplementary Figure 3: SynB1-ELP-p50i treatment does not reduce the increased expression of inflammatory cytokines, proteases, and adhesion molecules measured in brain lysate from the ipsilateral (red) hemisphere.** MCAO induced expression of all markers in the ipsilateral (red) hemisphere (two-way ANOVA; $F(3, 40) = 0.6380$, Šidák's post-hoc, $p < 0.05$). SynB1-ELP-p50i did not significantly reduce expression of any of these markers compared to the MCAO + Saline group. However, there was a clear distinction in expression of these markers between the contralateral (blue) and ipsilateral hemispheres, and there may be a downward trend in expression of ICAM-1, MMP-9, IL-6, and CXCL2 following MCAO + SynB1-ELP-p50i.

**Biodistribution of SynB1-ELP-p50i following IP Administration**
Intraperitoneal injection of rhodamine-labeled SynB1-ELP-p50i following 120 minutes of MCAO led to high deposition of SynB1-ELP-p50i in the brain, specifically in the infarcted core and penumbra, following 4 and 24 hours. Using this route of administration, a longer time was required, but equivalent deposition in the infarct to that observed after IV or IA injection was achieved within 24 hours post-injection.

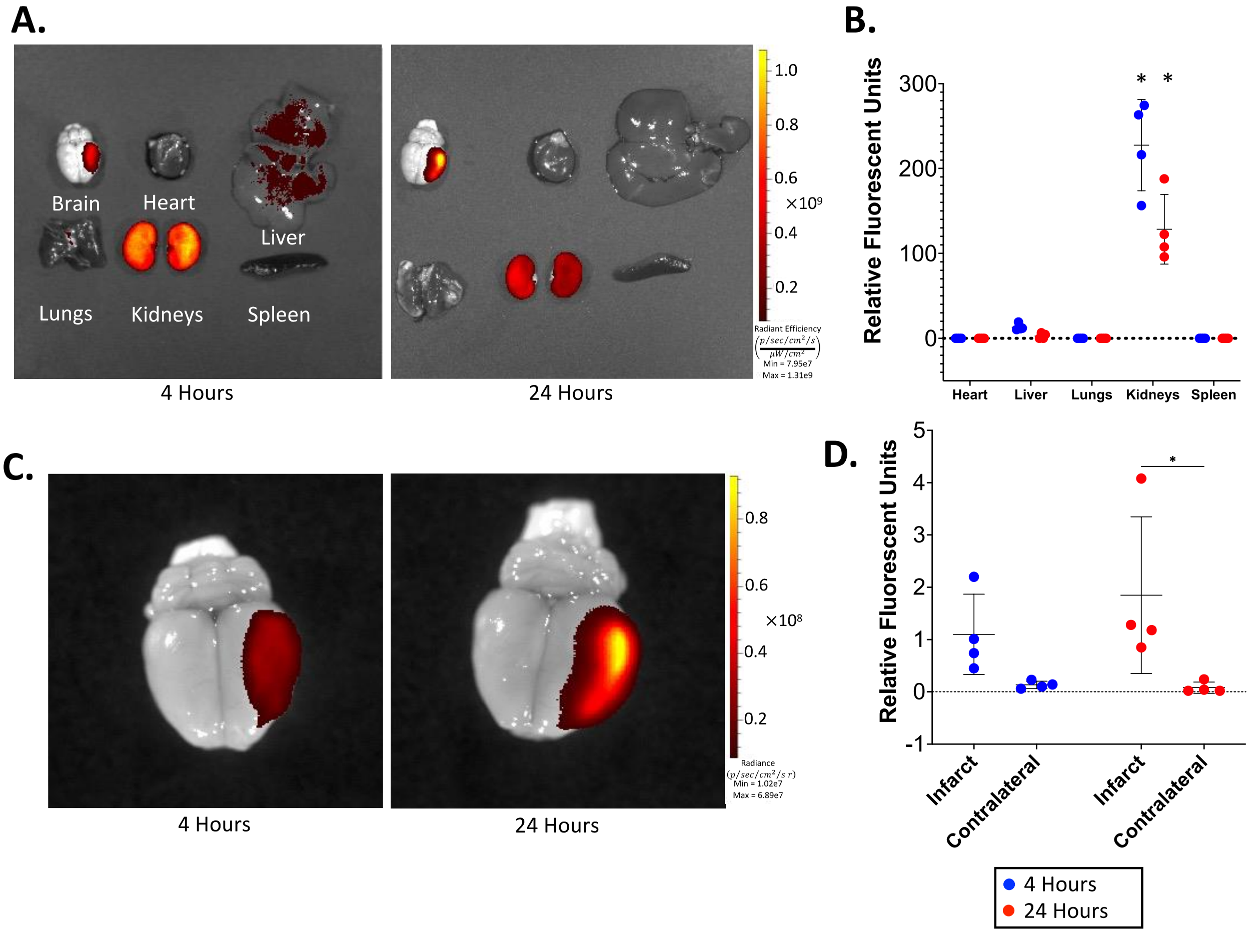


**Supplementary Figure 4: SynB1-ELP-p50i localizes to the infarct following intraperitoneal injection after MCAO in SHRs. A.** Representative IVIS images of IP SynB1-ELP-p50i treated SHRs at 4 hours and 24 hours post-injection. **B.** Quantified relative fluorescence of IP SynB1-ELP-p50i treated SHRs showed high accumulation in the kidneys compared to every other organ (two-way ANOVA; $F(4,30) = 108.1$, Šidák's post-hoc, $p < 0.0001$). **C.** Representative images of brains of IP SynB1-ELP-p50i treated SHRs at 4 hours and 24 hours post-injection in smaller field of view. **D.** Quantified relative fluorescence of brain deposition of IP SynB1-ELP-p50i treated SHRs at 4 hours and 24 showed significant increase of rhodamine labeled SynB1-ELP-p50i in the infarcted hemisphere following IP delivery at 24 hours post-injection (two-way ANOVA; $F(1,12) = 10.48$, Šidák's post-hoc, $p < 0.05$).

**Behavioral Assessment of Motor Function after Stroke and SynB1-ELP-p50i Treatment**
Behavior tests were conducted on days 1, 2, 3, 7, and 14 post-surgery to assess neurological and sensorimotor deficits in animals following sham or MCAO and treatment with saline or SynB1-ELP-p50i. Following 120 minutes of MCAO, all rats, regardless of treatment group, showed neurological deficit as measured by the Bederson score, decreased latency to fall on the rotarod, and increased time to remove adhesive stickers from forepaws. There were no differences among groups in performance in the open field test.

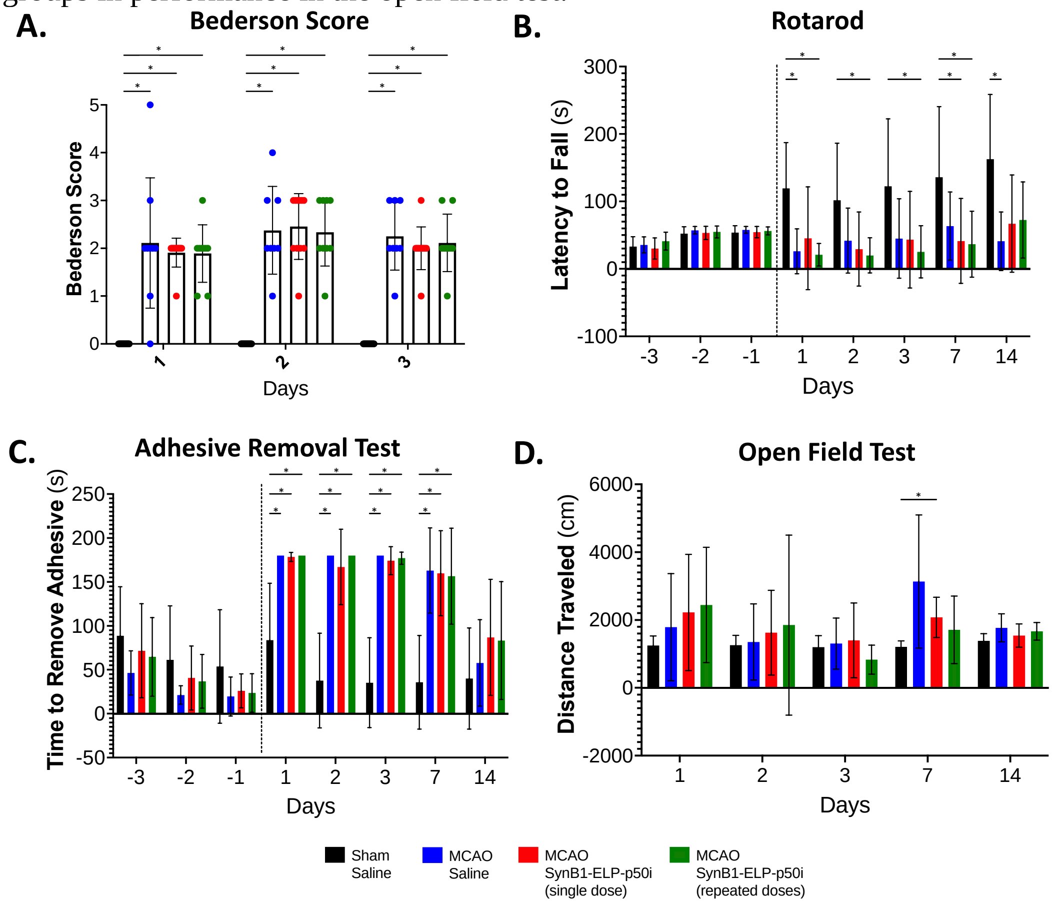


**Supplementary Figure 5: SynB1-ELP-p50i treatment has no effect on behavior outcomes following 120 minutes of MCAO. A.** Bederson score of neurological deficits showed significant impairment induced by MCAO (mixed-effects model (REML), Tukey's multiple comparisons test; $F(3, 40) = 44.47$, $p < 0.0001$) but no differences between Saline and SynB1-ELP-p50i treated MCAO groups on post-surgery days 1, 2, and 3. **B.** Results from accelerating rotarod assessments showed significant decreases in latency to fall in MCAO groups on days 1, 2, 3, 7, and 14 compared to the Sham group (mixed-effects model (REML), Tukey's multiple comparisons test; $F(3, 40) = 8.381$, $p < 0.05$). Dotted line denotes day of MCAO, and negative days indicate training days with a maximum time of 60 seconds. **C.** Time to remove adhesive was significantly increased in animals following MCAO surgery compared to Sham + Saline animals (mixed-effects model (REML), Tukey's multiple comparisons test; $F(3, 40) = 10.42$, $p < 0.05$). **D.** Total distance traveled during 10 minutes of open field test showed no significant differences between groups on days 1, 2, 3, and 14, but an increase in distance traveled by the MCAO + SynB1-ELP-p50i single dose group compared to Sham + Saline on day 7 (mixed-effects model (REML), Tukey's multiple comparisons test; $F(3, 40) = 1.852$, $p < 0.05$).

**Plasma Toxicology Markers 14 Days following Stroke and SynB1-ELP-p50i Treatment**
No differences were observed in markers of liver damage (ALT, AST), kidney damage (CREAT, BUN), or organ damage (LDH) at 14 days following sham or MCAO surgery, indicating that MCAO and SynB1-ELP-p50i treatment does not cause any toxicity when given as a single dose or repeatedly for 14 days.

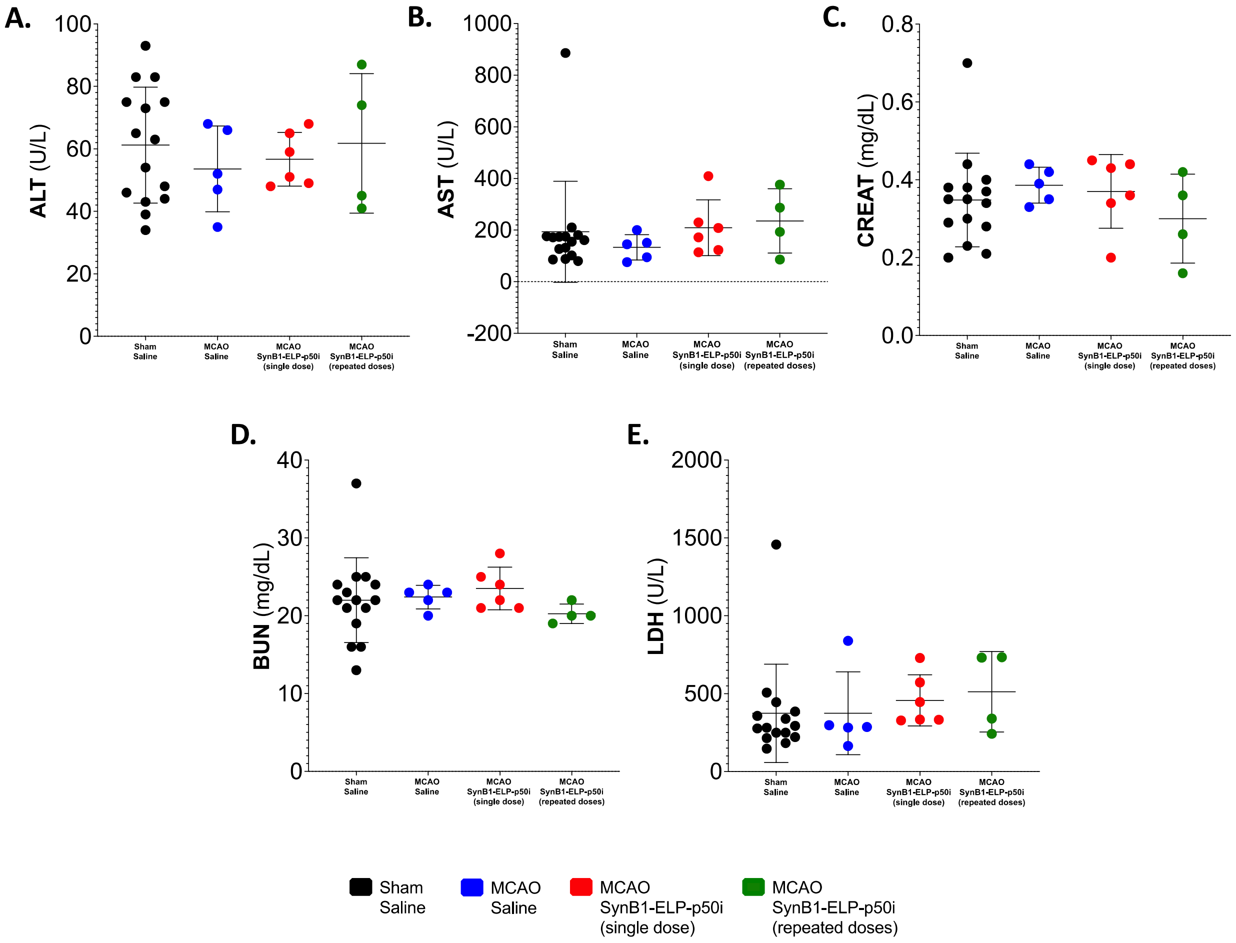


**Supplementary Figure 6: Treatment with SynB1-ELP-p50i following MCAO did not affect markers of liver damage (ALT, AST), kidney damage (CREAT, BUN), or organ damage (LDH) 14 days post-MCAO. A-E.** One-way ANOVA revealed no significant differences in levels of ALT ($F(3,26) = 0.3275$, $p = 0.8055$), AST ($F(3, 26) = 0.3541$, $p = 0.7866$), CREAT ($F(3, 26) = 0.5510$, $p = 0.6520$), BUN ($F(3, 26) = 0.4824$, $p = 0.6974$), or LDH ($F(3, 26) = 0.3499$, $p = 0.7895$).

**Chronic Effects of SynB1-ELP-p50i on Renal Fibrosis and Spleen Weight**
There were no differences among groups in kidney fibrosis or spleen weight at 14 days following sham or MCAO surgery, regardless of treatment condition.

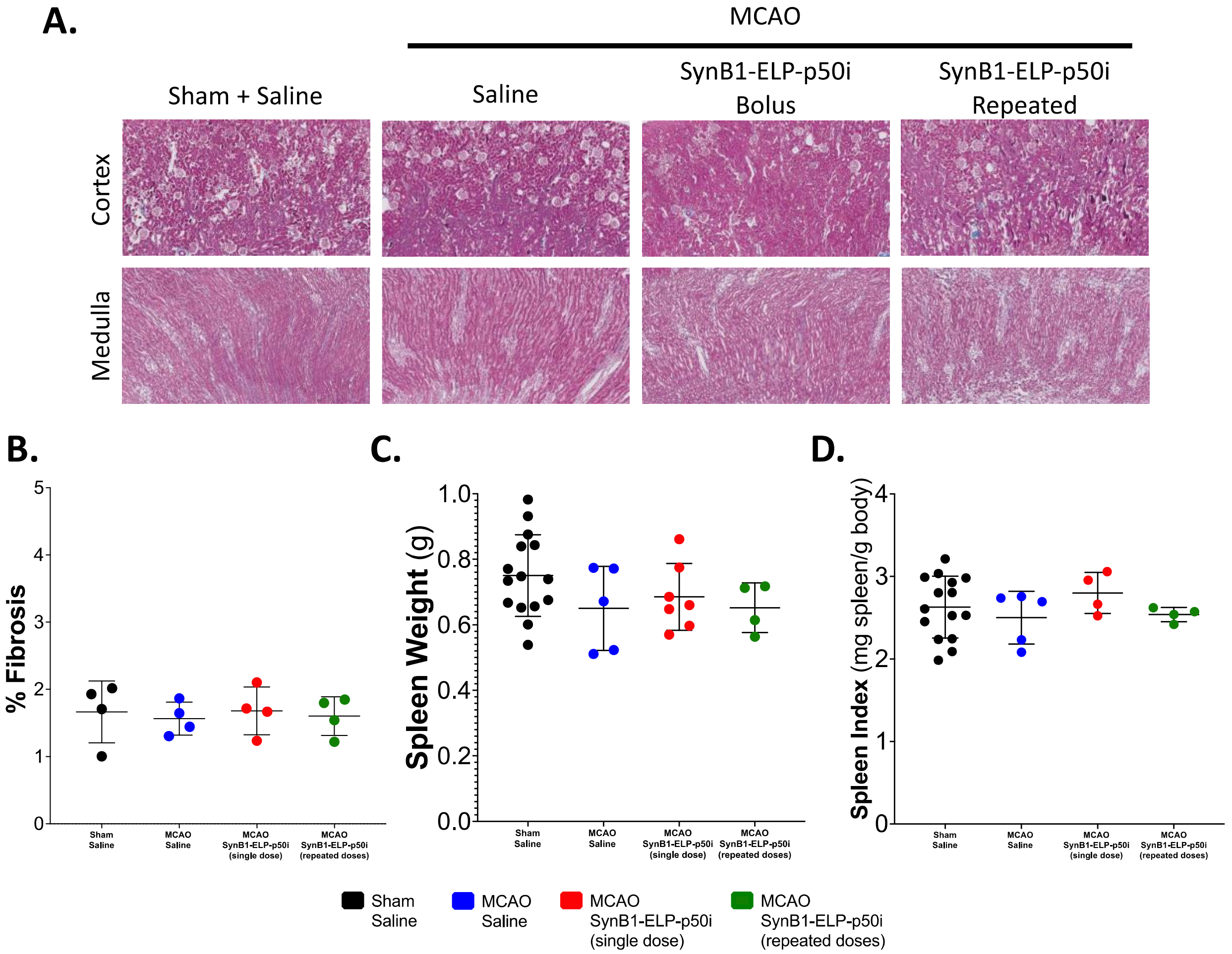


**Supplementary Figure 7: No kidney fibrosis or difference in spleen weight was seen following MCAO and treatment with saline or SynB1-ELP-p50i. A.** Representative Masson's trichrome stained images are shown from the renal cortex and medulla for each group. Collagen is stained blue in these images and is representative of fibrosis. Dark red shows cytoplasm or erythrocytes. **B.** There were no differences between groups in % Fibrosis for the entire kidneys (one-way ANOVA; $F(3,12) = 0.09578$, $p = 0.9609$). **C.** There were no differences between groups (one-way ANOVA; $F(3, 27) = 1.482$, $p = 0.2416$). **D.** To correct for any differences in body weight, spleen index was calculated by dividing the spleen weight (mg) by the animal body weight (g) at day 14. There were no differences between groups (one-way ANOVA; $F(3, 26) = 0.7101$, $p = 0.5555$).
